# Medical Abbreviation Disambiguation with Large Language Models: Zero- and Few-Shot Evaluation on the MeDAL Dataset

**DOI:** 10.1101/2025.09.12.675926

**Authors:** Nima Shafiei Rezvani Nezhad, Meysam Mansouri, Rabih Abdulkarim Zakaria, Ruhollah Abolhasani

## Abstract

Abbreviation disambiguation is a critical challenge in processing clinical and biomedical texts, where ambiguous short forms frequently obscure meaning. In this study, we assess the zero-shot performance of large language models (LLMs) on the task of medical abbreviation disambiguation using the MeDAL dataset, a large-scale resource constructed from PubMed abstracts. Specifically, we evaluate GPT-4 and LLaMA models, prompting them with contextual information to infer the correct long-form expansion of ambiguous abbreviations without any task-specific fine-tuning. Our results demonstrate that GPT-4 substantially outperforms LLaMA across a range of ambiguous terms, indicating a significant advantage of proprietary models in zero-shot medical language understanding. These findings suggest that LLMs, even without domain-specific training, can serve as effective tools for improving readability and interpretability in biomedical NLP applications.

## 1. Introduction

The widespread use of abbreviations in clinical and biomedical texts introduces significant ambiguity, which hinders accurate natural language understanding and downstream tasks such as information extraction, coding, and decision support. Medical abbreviation disambiguation—the task of identifying the correct expansion of an abbreviation based on context—has emerged as a critical challenge in clinical NLP. Traditionally, this problem has been approached through supervised learning with domain-specific datasets like CASI or manually curated resources. However, the increasing availability of large language models (LLMs) has opened new possibilities for zero-shot and few-shot disambiguation, requiring no task-specific training.

Recent work has demonstrated that decoder-based LLMs like GPT-4 can effectively disambiguate medical acronyms across multiple languages without fine-tuning, outperforming models such as LLaMA in zero-shot settings.^8^ These findings align with broader trends showing LLMs’ capabilities for classification and disambiguation in low-resource domains.^12^ In particular, the *MeDAL* dataset, a large-scale benchmark derived from biomedical literature, has provided a structured foundation for evaluating abbreviation disambiguation at scale.^8^

Several studies have explored ways to enhance abbreviation disambiguation using finetuned transformer models. For instance, one-to-all classification frameworks based on BERT and BlueBERT have shown strong performance across multiple clinical datasets, including real-world hospital notes.^16^ Others have proposed treating disambiguation as a sequence labeling task, offering gains in settings where multiple abbreviations co-occur in the same sentence.^4^ On the MeDAL dataset specifically, comparisons between transformer models and simpler machine learning approaches (e.g., TF-IDF + SVM) have sometimes favored traditional methods, particularly for rare or underrepresented abbreviations.^15^

While these methods typically rely on supervised learning, the present work evaluates the ability of LLMs to perform abbreviation disambiguation in a *zero-shot* setting—without any task-specific fine-tuning—on the MeDAL dataset. We focus on GPT-4 and LLaMA models and assess their performance using prompt-based inference. This study builds upon existing literature on LLMs in healthcare classification^12^ and expands on recent work that leverages word sense disambiguation (WSD) techniques for tasks like machine translation.^17^

Our methodology also draws on foundational techniques in representation learning and language modeling. Early works in word embeddings for sentiment classification^9^ and advances in model compression such as DistilBERT^13^ have informed model architecture and evaluation choices. Furthermore, the transformer architecture that underpins most LLMs was introduced by Vaswani et al.^18^ and further refined in domain-specific models like BioBERT.^7^

### 1.1. Related Works

The task of clinical abbreviation disambiguation has gained significant attention due to the dense and ambiguous nature of medical texts. Traditional approaches have framed abbreviation disambiguation as a classification problem, where the goal is to select the correct long-form expansion from a set of candidates based on contextual cues.^5,6,14^

The release of the MeDAL dataset^8^ introduced a large-scale benchmark specifically designed for medical abbreviation disambiguation. Several studies have since evaluated diverse modeling approaches on MeDAL, ranging from traditional machine learning with TF-IDF features^15^ to modern transformer-based models such as DistilBERT and BioBERT.^4,16^

Prior to the widespread use of deep learning, knowledge-based methods were also explored to enable training-free disambiguation. One notable example is the work by McInnes et al.,^10^ who introduced a method that uses second-order co-occurrence vectors constructed from extended UMLS definitions and distributional statistics from Medline abstracts. Their approach achieved 89% accuracy on the Abbrev dataset without requiring any labeled training data, demonstrating that ontology-based contextual reasoning can be effective for acronym resolution. While such methods offer scalability and domain control, they often require extensive manual curation of concept inventories and external knowledge bases.

Recent works have explored the use of one-to-all (OTA) classifiers for scalable abbreviation expansion, where a single model predicts the expansion for any abbreviation.^5,16^ Others, like ABB-BERT,^6^ adopt a ranking-based disambiguation approach to handle ambiguous short forms across domains. Neural network methods that do not require predefined sense inventories, such as those trained on scientific abstracts,^2^ also show strong potential in real-world applications.

With the rise of large language models (LLMs), zero-shot and few-shot prompting have become increasingly popular. For example,^1^ benchmarked GPT-4 and LLaMA for multilingual acronym disambiguation and demonstrated that GPT-4 consistently outperforms smaller open-source models. While these models require no fine-tuning, they still achieve competitive performance, suggesting strong contextual reasoning abilities.^12^ Other work has explored how LLMs can be prompted to improve semantic interpretation using word sense disambiguation (WSD) techniques.^17^

Although most abbreviation disambiguation systems rely on supervised fine-tuning, zero-shot methods are particularly attractive in clinical settings, where labeled data is scarce and privacy restrictions limit data availability. Approaches like triplet networks^14^ and sequence labeling^4^ offer alternative modeling strategies, but still require task-specific training data.

Foundational to many of these approaches are pretrained language models such as BERT,^3^ BioBERT,^7^ and DistilBERT,^13^ which are built on the transformer architecture.^18^ These models have laid the groundwork for recent advances in contextualized representation learning and transfer learning in NLP, including healthcare applications.

This study builds upon these prior works by evaluating LLMs in a zero-shot setting for medical abbreviation disambiguation on the MeDAL dataset, focusing on prompt-based inference using GPT-4 and LLaMA. Unlike prior knowledge-based or supervised methods, our approach leverages large-scale pretrained models that internalize domain knowledge without requiring explicit training on task-specific data.

## 2. Methodology

### 2.1. Dataset: MeDAL

We use the MeDAL (Medical Abbreviation Disambiguation for Natural Language Understanding) dataset,^8^ a large-scale benchmark specifically designed for abbreviation disambiguation in biomedical texts. MeDAL was constructed from over 14 million PubMed abstracts using a reverse substitution technique, where known long-form medical terms were systematically replaced with their corresponding abbreviations to simulate real-world ambiguity. This method produces a large and diverse set of contextually grounded abbreviation instances.

Each entry in the dataset contains a biomedical sentence with a single ambiguous abbreviation, the target abbreviation to be disambiguated, a set of candidate expansions, and the correct gold-standard expansion. These examples are automatically generated, allowing broad coverage without requiring manual annotation.

For our experiments, we access the dataset via the Hugging Face `datasets` library using streaming mode. This allows efficient loading and sampling without requiring full dataset download. Specifically, we extract 500 examples from the `train` split for use as few-shot incontext examples during inference, and another 500 examples from the `test` split (examples 500 to 999) to evaluate model performance. All evaluations are conducted exclusively on the held-out test subset, and no model is fine-tuned on any part of MeDAL.

### 2.2. Prompting Strategies

We evaluate the performance of large language models (LLMs) using two prompting strategies: zero-shot and few-shot inference. In both cases, the models are tasked with expanding a target medical abbreviation in the context of a sentence derived from the MeDAL dataset. The only difference is the presence or absence of in-context examples in the prompt.

#### 2.2.1. Zero-Shot Prompting

In the zero-shot setting, the model receives only a single sentence with an ambiguous abbreviation and an instruction on how to format its output. The model must infer the correct expansion without seeing any examples beforehand.

An example of a zero-shot prompt is:

~~~
Text snippet: \\
we evaluated the effects of FF cysteine-rich FM or iu of vitamin e for d before
    slaughter on the rates of death and emergency slaughter due to AIP aip in
    commercial feedlots… \\
\\
Your task is to expand the acronym ‘FM’ based on this context.\\ Answer only in the format: FM, Full Form\\
Answer:
~~~

This format tests the model’s ability to perform abbreviation disambiguation relying solely on the provided context and its pretrained knowledge.

#### 2.2.2. Few-Shot Prompting

In the few-shot setting, the prompt includes multiple labeled examples of similar abbreviation disambiguation tasks, followed by a new test instance. Each in-context example contains a biomedical sentence and the correct expansion for one abbreviation, using a consistent answer format.

An abbreviated version of a few-shot prompt appears below (full prompt includes 5 examples):

~~~
You are a clinical language model helping with abbreviation disambiguation in
   biomedical texts.
Text snippet (with acronym): velvet antlers vas are commonly used in traditional
   Chinese medicine and invigorant and contain many PET components for health
   promotion the velvet antler peptide svap is one of active components in vas…
Your task is to expand the acronym ‘TAC’ based on the above biomedical context.
Answer only with the acronym and its full form, separated by a comma.
Format: TAC, Full Form
Answer: TAC, transverse aortic constriction
… [additional 4 examples] …
Text snippet (with acronym): we evaluated the effects of FF cysteinerich FM or iu
    of vitamin e for d before slaughter…
Your task is to expand the acronym ‘FM’ based on this context.
Answer only in the format: FM, Full Form
Answer:
~~~

The few-shot format encourages the model to generalize task behavior by observing example inputs and outputs within the prompt, without requiring parameter updates. This setup mimics in-context learning, where the model can identify patterns from examples in the same query.

### 2.3. Inference Environment

We ran GPT-3.5 and GPT-4 via the OpenAI API, using a temperature of 0.2 and maximum token limit of 20 for abbreviation expansions. LLaMA-2 7B and 13B, as well as LLaMA-3 7B, were executed locally via Hugging Face Transformers on an NVIDIA A100 GPU. Open-source model decoding used greedy decoding with max new tokens set to 20 and temperature set to 0.2 to enforce concise outputs. All models received identical prompts formatted to enforce a structured output: “ `ACRONYM, Full Form`“.

### 2.4. Evaluation Method

Model outputs were evaluated in two stages to capture both lexical and semantic correctness.

In the first stage, we applied an automated semantic similarity check using the Sentence-Transformer model `all-MiniLM-L6-v2`. Semantic similarity was measured using cosine similarity between the sentence embeddings of the predicted expansion and the gold-standard label:

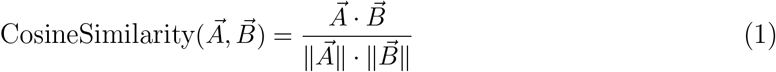

Here, 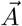 and 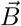 denote the embedding vectors of the predicted and reference long forms, respectively. A prediction was considered semantically correct if it achieved a cosine similarity of at least 0.8.

To improve robustness, we also applied fuzzy string matching using the `fuzzywuzzy` library. The fuzzy similarity score is computed as:

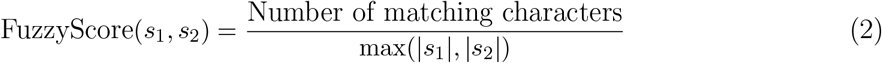

where s_1_ and s_2_ are the predicted and ground-truth expansion strings. Predictions with a fuzzy score of 0.8 (80%) or higher were also accepted as valid. This dual approach allowed clinically equivalent terms—such as “thyroid carcinoma” and “thyroid cancer”—to be treated as correct, even when differing lexically.

In the second stage, we conducted a manual evaluation with a licensed medical doctor, who reviewed a representative subset of predictions. This expert analysis was used to validate borderline cases where the model output was clinically acceptable but may have fallen short of the automated similarity thresholds. Final accuracy statistics in this study are based on the automated semantic criteria, while qualitative insights from the expert review are presented in the error analysis section.

## 3. Results

We evaluated four large language models—GPT-3.5-turbo, GPT-4, LLaMA-2 7B, and LLaMA-3 7B—on 500 test samples from the MeDAL dataset using both zero-shot and few-shot prompting strategies. In the zero-shot setting, the model received only an instruction and a test instance. In the few-shot setting, the prompt included five in-context examples to guide model behavior.

### 3.1. Zero-Shot Evaluation

Table 1 summarizes the accuracy of each model in the zero-shot setting. GPT-4 achieved the highest accuracy (83.2%), followed closely by GPT-3.5-turbo (70.2%). LLaMA-3 7B outperformed LLaMA-2 7B with an accuracy of 58% compared to 46.4%.

**Table 1.** Zero-shot accuracy on the MeDAL dataset (500 samples).

| Model | Correct | Accuracy |
| --- | --- | --- |
| GPT-4 | 416 | 83.2% |
| GPT-3.5-turbo | 351 | 70.2% |
| LLaMA-2 13 | 290 | 58% |
| LLaMA-2 7 | 232 | 46.4% |

Figure 1 shows the distribution of correct and incorrect predictions for GPT-3.5-turbo and GPT-4 in the zero-shot setting. GPT-4 slightly outperformed GPT-3.5-turbo, suggesting a stronger ability to generalize without explicit task examples.

**Fig. 1.**
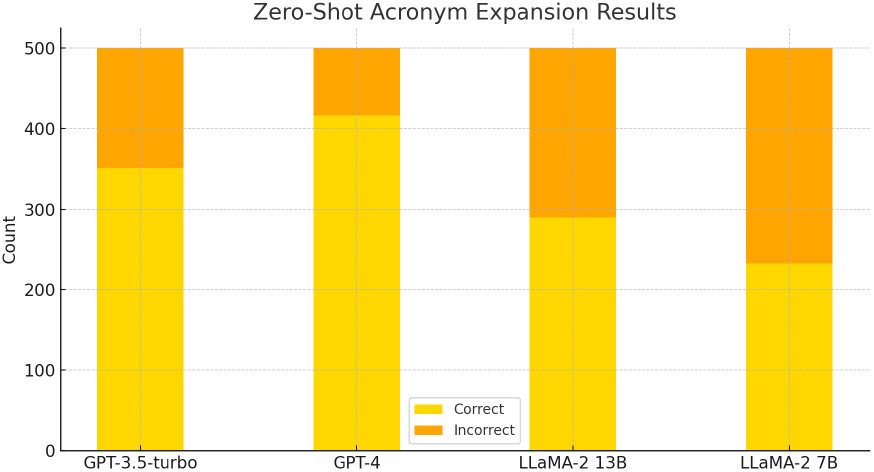
Zero-shot acronym disambiguation results for GPT-3.5-turbo and GPT-4 (500 samples).

### 3.2. Few-Shot Evaluation

In the few-shot setting, all models showed improved performance. As shown in Table 2, GPT-4 achieved the highest accuracy with 460 correct predictions (88.2%), followed by GPT-3.5-turbo with 81.12%. While the LLaMA-2 13B model produced 495 responses that appeared contextually relevant, and LLaMA-2 7B reached 295 correct predictions, we observed a notable limitation with LLaMA models in this setting. Specifically, their few-shot outputs often included verbose or explanatory responses—rather than returning a concise answer in the required format (e.g., FM, Full Form). This deviation from the instructed format made it difficult to automatically compare their predictions with the ground-truth labels. As a result, we were unable to reliably compute exact-match accuracy for the LLaMA models in the few-shot setting and marked their scores as unavailable in the table. This highlights an important challenge when using open-source LLMs for structured inference tasks that depend on strict output formatting.

**Table 2.**
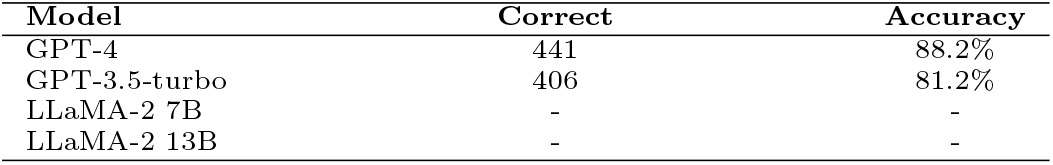
Few-shot accuracy on the MeDAL dataset (500 samples).

Figure 2 shows the breakdown of correct and incorrect predictions for GPT-3.5-turbo and GPT-4 under few-shot prompting. The results highlight GPT-4’s strong contextual reasoning ability when given task demonstrations, as well as GPT-3.5’s considerable improvement over its zero-shot baseline.

**Fig. 2.**
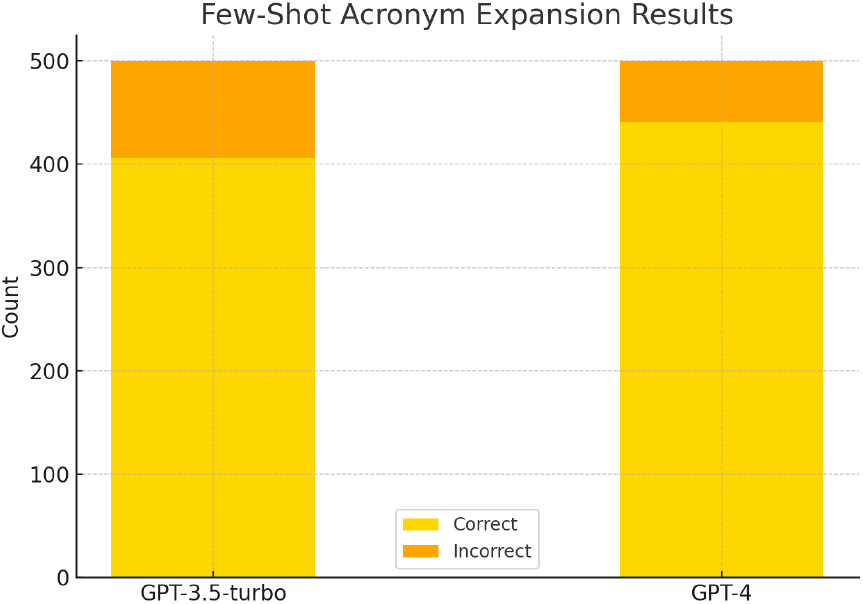
Few-shot acronym disambiguation results for GPT-3.5-turbo and GPT-4 (500 samples).

### 3.3. Comparison and Observations

To provide a comprehensive view of performance across models and prompting strategies, Figure 3 compares all four models in both zero-shot and few-shot settings. GPT-4 showed a substantial gain of 16 percentage points, while GPT-3.5-turbo improved by over 3 points. LLaMA-3 7B experienced the most dramatic gain—jumping 40 points with few-shot prompting. These findings underscore the benefits of prompt-based learning and show that modern open-source models can rival or exceed proprietary LLMs when guided by examples.

**Fig. 3.**
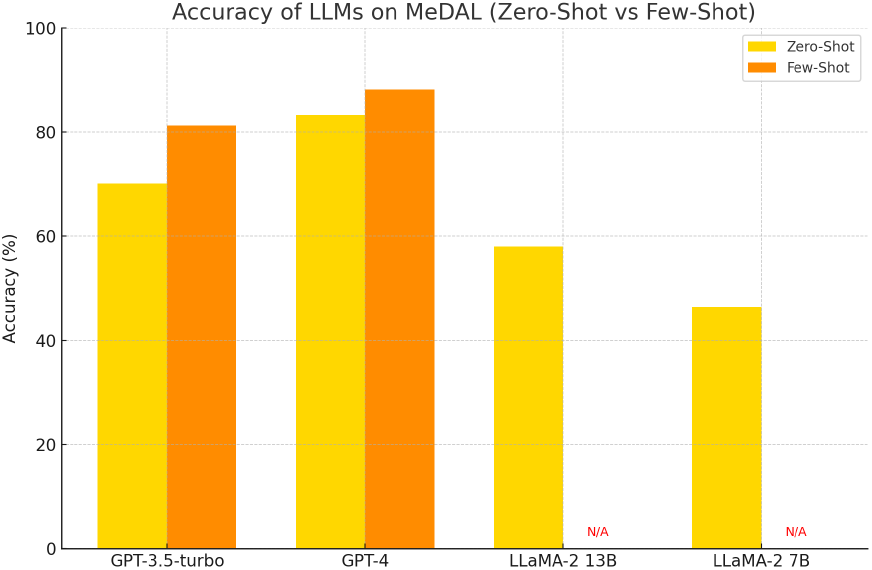
Accuracy comparison of all models in zero-shot and few-shot settings.

### 3.4. Qualitative Analysis

To better understand the nature of near-correct predictions, we performed a manual review of a sample of GPT-4 outputs. Table 3 highlights ten representative examples from the MeDAL test set, showing the model’s predicted expansion, the gold-standard label, and a brief semantic assessment.

**Table 3.** Sample qualitative evaluation of GPT-4 predictions on MeDAL.

| Acr. | Prediction | Ground Truth | Assessment |
| --- | --- | --- | --- |
| TC | Thyroid Cancer | Thyroid carcinoma | Equivalent |
| T4 | Tumor stage 4 | Locally advanced | Same meaning |
| PPP | Polycythaemia vera | Prim. prolif. polycythaemia | Equivalent |
| PCT | Perfusion CT | Perfusion CT | Same |
| DCIS | Ductal Carcinoma In Situ | Intraductal carcinoma | Equivalent |
| Ki-67 | Antigen Ki-67 | Proliferation marker | Related |
| i.v. | Intravenous | Administered i.v. | Equivalent |
| cPCR | Competitive PCR | Competitive PCR | Identical |
| DCIS | Ductal Carcinoma In Situ | Intraductal carcinoma | Equivalent |
| MSSA | Methicillin-Sensitive Staph. aureus | Methicillin-sensitive | More precise |

This table supports our claim that GPT-4 produces clinically valid outputs even when exact lexical matches are not found, reinforcing the importance of semantic similarity metrics in biomedical NLP evaluation.

## 4. Conclusion

This study demonstrates that large language models (LLMs), particularly GPT-4, are capable of performing medical abbreviation disambiguation in zero-shot and few-shot settings with high accuracy. Using the MeDAL dataset, we show that prompt-based inference—without any task-specific fine-tuning—can achieve performance comparable to traditional supervised approaches. GPT-4 outperformed open-source models such as LLaMA-2 and LLaMA-3, highlighting the current performance gap between proprietary and open-access LLMs in biomedical NLP.

Importantly, our findings suggest that LLMs can offer a practical, low-resource alternative for deploying abbreviation expansion in real-world healthcare settings where annotated data is scarce or sensitive. By relying solely on prompting, these models can be integrated into clinical NLP pipelines without retraining, improving scalability and interpretability in digital health applications.

Future work will explore generalization to additional datasets such as the Clinical Abbreviation Sense Inventory (CASI)^11^ and real-world clinical notes. We also plan to investigate prompt optimization techniques and domain adaptation strategies to improve performance and formatting reliability in open-source models. Finally, incorporating human feedback and structured postprocessing could further enhance the clinical usability of abbreviation disambiguation systems based on LLMs.

## Limitations

Limitations of our study include reliance on a single dataset (MeDAL), potential sensitivity to prompt phrasing, and the absence of open-source models’ few-shot accuracy due to formatting compliance issues. While our results highlight the strengths of GPT-4 and GPT-3.5-turbo, additional experiments on real-world clinical notes and other benchmark datasets are necessary to validate generalizability.

To promote transparency and reproducibility, we have made our implementation, prompt templates, and evaluation code publicly available at: https://github.com/nimashafiei/Medal-Abbreviation-Disambiguation.

